# Single cell atlas of domestic pig brain illuminates the conservation and divergence of cell types at spatial and species levels

**DOI:** 10.1101/2019.12.11.872721

**Authors:** Dongsheng Chen, Jiacheng Zhu, Jixing Zhong, Fang Chen, Xiumei Lin, Jinxia Dai, Yin Chen, Shiyou Wang, Xiangning Ding, Haoyu Wang, Jiaying Qiu, Feiyue Wang, Weiying Wu, Ping Liu, Gen Tang, Xin Qiu, Yetian Ruan, Jiankang Li, Shida Zhu, Xun Xu, Fang Li, Zhongmin Liu, Gang Cao

**Affiliations:** BGI-shenzhen, Shenzhen 518083, China; BGI Education Center, University of Chinese Academy of Sciences, Shenzhen 518083, China; School of Biological Science & Medical Engineering, Southeast University, Nanjing 210096, China; State Key Laboratory of Agricultural Microbiology, Huazhong Agricultural University, Wuhan 430070, China; College of Veterinary Medicine, Huazhong Agricultural University, Wuhan 430070, China; Key Laboratory of Development of Veterinary Diagnostic Products, Ministry of Agriculture, College of Veterinary Medicine, Huazhong Agricultural University, Wuhan 430070, China; School of Life Sciences, South China Normal University, Guangzhou 510631, China; MGI, BGI-Shenzhen, Shenzhen 518083, China; Shenzhen Children’s Hospital, Shenzhen 518083,China; Research Center for Translational Medicine, East Hospital, Tongji University School of Medicine, 150 Jimo Road, Shanghai 200120, China

## Abstract

Domestic pig (*Sus scrofa domesticus*) has drawn much attention from researchers worldwide due to its implications in evolutionary biology, regenerative medicine and agriculture. The brain atlas of *Homo sapiens* (primate), *Mus musculus* (rodent), *Danio rerio* (fish) and *Drosophila melanogaster* (insect) have been constructed at single cell resolution, however, the cellular compositions of pig brain remain largely unexplored. In this study, we investigated the single-cell transcriptomic profiles of five distinct regions of domestic pig brain, from which we identified 21 clusters corresponding to six major cell types, characterized by unique spectrum of gene expression. By spatial comparison, we identified cell types enriched or depleted in certain brain regions. Inter-species comparison revealed cell-type similarities and divergences in hypothalamus between mouse and pig, providing invaluable resources for the evolutionary exploration of brain functions at single cell level. Besides, our study revealed cell types and molecular pathways closely associated with several diseases (obesity, anorexia, bulimia, epilepsy, intellectual disability, and autism spectrum disorder), bridging the gap between gene mutations and pathological phenotypes, which might be of great use to the development precise therapies against neural system disorders. Taken together, we reported, so far as we know, the first single cell brain atlas of *Sus scrofa domesticus*, followed by comprehensive comparisons across brain region and species, which could throw light upon future evo-devo, regenerative medicine, and agricultural studies.

## Introduction

The domestic pig (*Sus scrofa domesticus*), one of the most important livestocks, shares close interaction relationships with humans during evolution history[1–3]. It belongs to the eutherian mammal from cetartiodactyla order, a clade distinct from primates and rodents[4]. Domestic pig has been widely studied because of its significances in evolutionary biology and regenerative medicine[5]. During the past decades, pig is increasingly used in neuroscience researches, due to the much higher correspondence in its brain to human brain in anatomy, physiology, and development than that of commonly used small laboratory animals such as mouse and rat[6,7]. Similar to human brain, pig brain is gyrencephalic and easy to recognize anatomically. For example, the cerebral cortex is clearly differentiated into four lobes including temporal lobe (TL), frontal lobe (FL), parital lobe (PL), and occipital lobe (OL)[8]. Recently, a transgenetic pig Huntington’s disease (HD) model was established by a mutant huntingtin knockin, who shows classical HD phenotypes such as behavioral abnormalities, consistent movement and early death[9]. Accumulating genetic manipulation tools have facilitated the applications of pig as an animal model for researches into brain function, particularly studies concerning brain malfunctioning mechanism. Besides, for ethical and economical reasons, pig is a more acceptable lab animal than primate. Studying the pig brain is of crucial importance to understand the evolution, function and disease pathology of the central nervous system. Historically, the domestication of pig can date back to early human civilization. Nowadays, pig has become a pivotal livestock and the main source of meat, as a result of which, it is drawing more and more attention to developing finer breeding programs [10,11]. Several genomic studies has been conducted to dissect the molecular mechanisms and cellular circuit associated with pig growth, diet and body weight regulation and fat accumulation[1,12–17].

The brain of vertebrates, composed of billions of cells, is an extremely complex organ with high heterogeneity. Brains from different species have been extensively investigated using a variety of methods[18–24]. Yet, bulk data is rather limited in revealing molecular signature within distinct cell types, especially rare cell populations such as stem cells or cells under intermediate status[25,26]. Little is known about the complicated structural configuration and functionalities of pig brain at single cell resolution, for instance, subtle neuron type classification and detailed intercellular crosstalks, narrowing their applications in functional and pathological studies.

Since 2009, single cell RNA sequencing (scRNA-seq) technique has been applied to a wide range of studies due to its unique capability to disentangle the heterogeneity of complex tissues and precious samples with limited numbers of cells[27]. Brain, characterized by remarkable cellular diversity and intricate interplays, naturally fell into the ideal object of scRNA-seq inverstigations. Heretofore, scRNA-seq has been utiliezed to construct single cell atlas of certain brain region or nucleus from several species including human, mouse, fruit fly and zebrafish[28–33]. For human, developing cerebral cortex was first delineated at single cell level using Microfluidic C1 in 2014[28]. Thereafter, a series of subsequent studies provided more detailed characterizations of human brain in regionally broader coverage (cerebral cortex, cerebellum, midbrain, and temporal lobe) [29–31]. Additionally, brains from patients under different pathological conditions, Alzheimer’s disease (AD), autism spectrum disorder (ASD), and Parkinson’s disease (PD) included, were also interrogated using scRNA-seq and extensive dysfunctions were observed at unprecedent resolution[32–34]. For mice, single cell transcriptome of cortex and hippocampus were reported by Zeisel et. al.[35]. After that, a variety of methods were applied to reconstruct cellular hierarchies and developmental trajectories of neural cells as well as interactions between neural cells and immune cells[36]. Brains of other model animals such as fruit fly and zebrafish were also under investigation using single cell methods. Briefly, olfactory projection neuron, aging brain, and midbrain were analyzed to understand neural cell subtypes[37]. In zebrafish, scRNA-seq was used to dissect cellular lineages and cell types in both brains from healthy and amyloid toxicity model[38].

Pig brain has been investigated in the apsects of genome, transcriptome and epigenome, establishing a basic preception structurally and functionally [24,39]. However, bulk data is intrinsically restricted in dissecting the heterogeneity and micro-environment of complex tissue. Considering the importance of pig brain for neuroscience study, it is of great necessity to elucidate the single cell atlas of it using scRNA-seq, particularly the key functional regions including cortex and hypothalamus. Leveraging the advantages of pig model, we aim to gain deeper understanding of human cognition functioning and also, to promote the refinement of current pig farming guidelines. Therefore, we, with the help of scRNA-seq, comprehensively delineated the cellular compostion and molecular circuits within cerebral cortex and hypothalamus, which is the center of cognition formation and hormonal regulation respectivly. In concrete terms, we generated transcriptomic profiles of individual cells from four lobes in cerebral cortex (temporal lobe (TL), frontal lobe (FL), parietal lobe (PL), and occipital lobe (OL)) and hypothalamus (HT). We identified 21 clusters of cells based on unsupervised clustering, and further revealed the existence of cellular interaction networks within pig brain. We also identified certain cell clusters that show significant correlations with certain nervous system diseases. By cross-species comparison, we revealed several important neural function genes may play critical roles in the pig neural system. Our study may enhance the understanding of pig brain fine structure at single cell level, providing a solid platform for further agricultural and pathological researches regarding the development of noval regenerative medicine, cost-efficient breeding strategies etc.

## Results

### Generation of pig brain atlas at single cell resolution

To comprehensilvely characterize pig brain at single cell level, we constructed single cell libraries for each brain regions under platform, followed by high throughput RNA sequencing. After filtering, we obtained the single-cell transcriptome of 32250 cells from five brain regions, with 6829, 8812, 6162, 3145 and 7302 cells from OL, FL, PL, TL and HT respectively (Figure 1a). Generally, 21 cellular clusters were identified and visualized on a UMAP plot (Figure 1b). We, based on the expression of cell type marker genes, assigned 21 clusters to six major cell types: excitatory neurons (EX), inhibitory neurons (IN), oligodendrocyte progenitor cells (OPC), oligodendrocytes (OLG), astrocytes (AST), and microglias (MG). Cluster 3, 4, 8, 11, 13, 14, 15 were annotated as EX because of the expression of *SATB2, TLE4*, and *CUX2* (Figure 1f). Cluster 10, 12, 16, 19 and 20 were considered as IN due to the expression of *GAD1*, *GAD2*, and *NXPH1* (Figure 1f). Cluster 2, 9 were considered to be OPC indicated by the preferential expression of *VCAN*, *LHFPL3*, and *PTPRZ1* (Figure 1f). Cluster 0, 6 were recognized as OLG for the expression of *MBP*, *ERBIN*, and *CNP* (Figure 1f). Cluster 7 was annotated as AST characterized by the expression of *GFAP*, *SLC1A2*, and *SLC1A3* (Figure 1f). Cluster 1 was annotated as MG because of the expression of *P2RY12*, *CALCR*, and *ARHGAP24* (Figure 1f). We also identified a set of functionally diverse genes in each cluster and consistent with annotated cell identity, the specific gene set found in each cluster converge to specific biological functions matched cell identities, further confirming the validity of the identification of cell types. For example, the cluster specific genes of EX and IN were enriched for GO terms like “synapse organization” and “axonogenesis”. While, for OPC, OLG, and AST, the cluster specific genes showed enrichment for molecular functions associated with “gliogenesis”. Specific genes of MG were enriched for GO terms related to “lymphocyte differentiation” and “regulation of myeloid cell differentiation”. Apart for this, different combinations of these molecular fingerprints can be employed for the sorting of specific clusters of interests (Table S1, Figure1d).

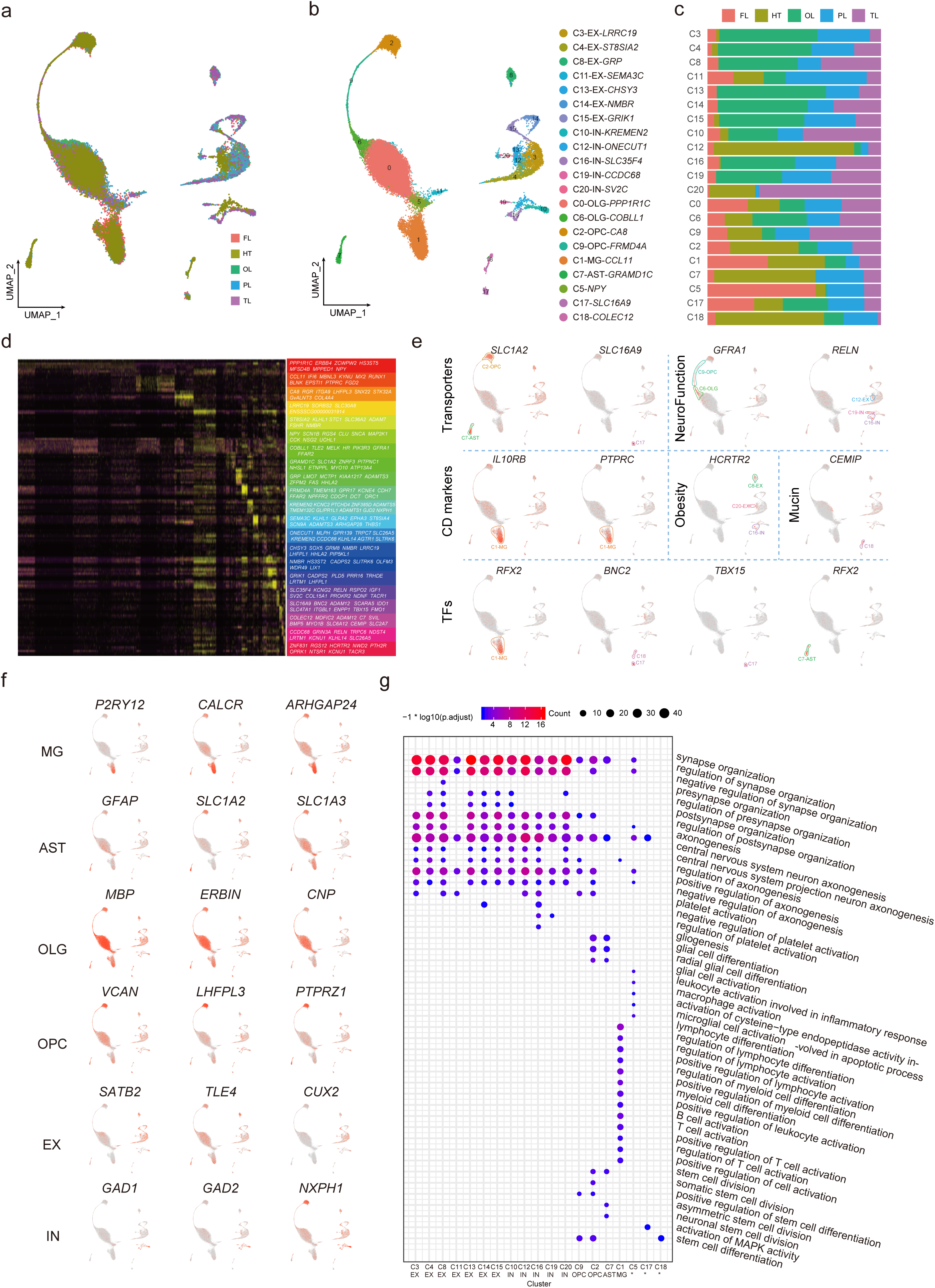

### Heterogeneity of transcription factors and functional genes within clusters

Transcription factors hold a pivotal role in molecular function activation by spatially and temporally regulating the precise expression patterns of certain gene modules. We then asked, to what extent, TFs expression varied among clusters. As expected, we identified a variety of cluster specific TFs. For example, C9 (OPC) specifically expressed 11 TF encoding genes (*ARNTL*, *TCF7L2*, *NPAS2*, *DACH1*, *SOX6*, *FOXP3*, *PBX3*, *SOX13*, *POU6F2*, *SOX5*, and *TOX* (Table S1)). TCF7L2 is essential for neurogenesis and has been reported to be associated with diabetes[40–42]. *ARNTL* is a well-known circadian clock gene[43,44]. NPAS2 regulates anxiety-like behavior and non-rapid eye movement sleep[45,46]. SOX6 is associated with the inhibition of neuronal differentiation[47]. TOX is proposed to be a versatile TF regulating mammalian corticogenesis[48].

C11 (EX) specifically expressed 16 TF encoding genes (*REV1*, *ZSCAN31*, *NEUROD2*, *ARID3B*, *TOX3*, *SATB2*, *ZNF831*, *ARNT2*, *MEOX1*, *POU3F2*, *PBX1*, *POU6F2*, *PRRX1*, *SOX5*, *ONECUT1*, and *SETBP1* (Table S1)). TOX3 was reported to regulate the neural progenitor identity, and was proposed to promote neuronal survival[49,50]. SATB2 regulates the identity of callosal projection neuron and contributes to the cognitive ability[51,52]. *RFX2* were specifically expressed in C7 (AST).

In addition to master regulators like TFs, we also scrutinized the expression of other functional genes (CD markers, transporter genes, glutamatergic associated genes, and GABAergic associated genes) that were also important for the biological functions of neural and glia cells [53] (Figure 1e). We noticed that two CD markers *IL10RB* and *PTPRC* were specifically expressed in C1 (MG). Transporter gene *SLC1A2* was specifically expressed in C2 (OPC) and C7 (AST). The neurological function gene *GFRA1* was specifically expressed in C9 (OPC) and C6 (OLG). *RELN*, also a neurological function gene, was specifically expressed in C12 (EX), C16 (IN) and C19 (IN). Interestingly, an obesity related gene *HCRTR2* was found to be specifically expressed in C8 (EX), C16 (IN) and C20 (IN).

Besides those well-defined clusters, we also found three clusters (C5, C17, C18), probably correspond to previously poorly characterized cell types. C5 preferentially expressed 27 genes including *SNAP25*, *NPY*, and *CCK* (Table S1). *SNAP25*, encoding a 25 kDa synaptosomal-associated protein, is related to several neurological diseases including schizophrenia[54] and attention-deficit/hyperactivity disorder (ADHD)[55]. *NPY* encodes a neuropeptide involved in multiple homeostatic and physiological processes of the nervous system[56,57]. *CCK* is a versatile molecular switch of neural circuits[58]. C17 specifically expressed 15 TF encoding genes (*TCF7L2*, *TBX15*, *ID3*, *NR3C2*, *AHR*, *NR2F1*, *SMAD3*, *PRDM6*, *PRDM5*, *THRB*, *KLF5*, *LIN28B*, *BNC2*, *ESR2*, and *KLF12*). SMAD3 co-activate neural developmental program with JMJD3 and regulates neuronal differentiation[59]. LIN28B regulates proliferation and neurogenesis in NPCs[60]. KLF12 is linked with cell proliferation[61,62]. C18 specifically expressed 19 TF encoding genes (*GTF2IRD1*, *GLI3*, *ETV1*, *ID3*, *SATB1*, *TEAD1*, *NPAS4*, *DACH1*, *AHR*, *ARID3B*, *ONECUT2*, *KLF5*, *MEOX1*, *BNC2*, *SOX13*, *TBX18*, *HIVEP1*, and *RORA*). Among them, GLI3 is required for the maintenaning and fate specifying of cortical progenitors, and it is proposed to regulate the migration of precerebellar neurons[63,64].

### Spatial orgins of each cluster

To measure the contribution of each brain region to every cell cluster, we calculated a customized metric named CI (contribution index), defined as the proportion of cell from a specific brain region in a given cell cluster, followed by normalization of the total number of cells from the brain region (Table S2). We used a intuitionistic criterion to measure the enrichment or depletion of certain brain regions in each cluster. Cluster with CI less than 10% or greater than 40% was considered to be depleted or enriched for a specific cell cluster. In this regrad, we noticed that EX subpopulations were mainly dominant by cells from OL (C3, C4, C8, C13, C14, C15), with one exception that C11 was dominant by cells from PL. We observed a distinct phenomenon in terms of IN where different IN subpopulations displayed a region-specifc distribution: C10 and C20 were enriched with cells from TL while C16 was prevailed by OL cells. Besides, HT showed predominance in C12. On the other hand, glial cells dispersed relatively equally across different brain regions and only C7 (AST) and C9 (OPC) exhibited enrichment with HT and TL respectively. We presumed that theses observations may indicate a conserved role of glial cells in cerebral cortex and hypothalamus. Nonetheless, neurons were comparatively more region-restricted, implying their contribution to the formation of the derivation of region-specific functions (Figure 1c, Table S2).

### Cellular heterogeneity and micro-environment in hypothalamus

The hypothalamus coordinates appetite control, energy balance through neuroendocrine circuits, essential for maintaining body homeostasis[65]. In total, 15 cellular clusters were identified in pig hypothalamus, corresponding to EX (C9, C14), IN (C4, C10, 12), MG (C1), OLG (C0, C2), OPC (C3, C6) and AST (C5, C7) (Figure 2a). CD markers (*ALK*, *BMPR1B*, *CD38*, *CD44*, *CDCP1*, *CDH2*, *FGFR1*, *ITGA6*, *KIT*, *MME*, *NRP1*, *SDC2*, *TFRC*, and *TNFRSF*) showed specific expression patterns in C3 (OPC), C4 (IN), C5 (AST), C6 (OPC), C7 (AST), C8, C9 (EX), C10 (IN), C11, C12 (IN), and C14 (EX) (Figure 2c). Appetite regulators (*CCKAR* and *HCRTR2)* were specifically expressed in C10 (IN) and C4 (IN) respectively. Alpha chain of type VIII collagen encoding gene *COL8A1* was specifically expressed in C5 (AST), C9 (EX), C10 (IN), and C11, while *COL4A2* was preferentially expressed in C9 (EX) and C11. Epinephrine receptor *ADRA1A* was expressed in C12 (IN). Dramamine receptor (*DRD2*) was expressed in C10 (IN) and dopamine receptor *SLC6A3* was specifically expressed in C10 (IN) and C13. GABA receptors (*GABBR2*, *GABRA1*, *GABRA2*, *GABRA3*, *GABRA5*, *GABRB1*, *GABRB2*, *GABRB3*, *GABRD*, and *GABRG2*) were mainly expressed in C3 (OPC), C4 (IN), C5 (AST), C7 (AST), C9 (EX), C10 (IN), C12 (IN), and C14 (EX). Glutamate receptors (*GRIA2*, *GRIA3*, *GRIA4*, *GRM5*, *GRIA1*, *GRIA2*, *GRIA3*, *GRIA4*, *GRM1*, *GRM4*, *GRM5*, *GRM8*, *GRM5*, *GRIA1*, *GRIA3*, *GRIA4*, *GRIA1*, *GRIA2*, *GRIA3*, *GRM1*, *GRM4*, *GRIA1*, *GRIA2*, *GRIA3*, *GRIA1*, *GRIA2*, *GRIA3*, *GRIA4*, *GRM1*, *GRM5*, *GRM8*, *GRIA1*, *GRIA2*, *GRIA3*, *GRM1*, *GRM5*, and *GRM8*) were mainly expressed among C3 (OPC), C4 (IN), C5 (AST), C6 (OPC), C9 (EX), C10 (IN), C12 (IN), and C14 (EX) (Figure 2c). Mucin (*CEMIP*) was mainly expressed in C8 (Figure 2c). Nicotinic acetylcholine receptors (*CHRNA4*, *CHRNA6*, and *CHRNB3*) were specifically expressed in C10 (IN), while *CHRNA7* was specifically expressed in C12 (IN). Obesity related gene *HCRTR2* was specifically expressed in C4 (IN) (Table S13).

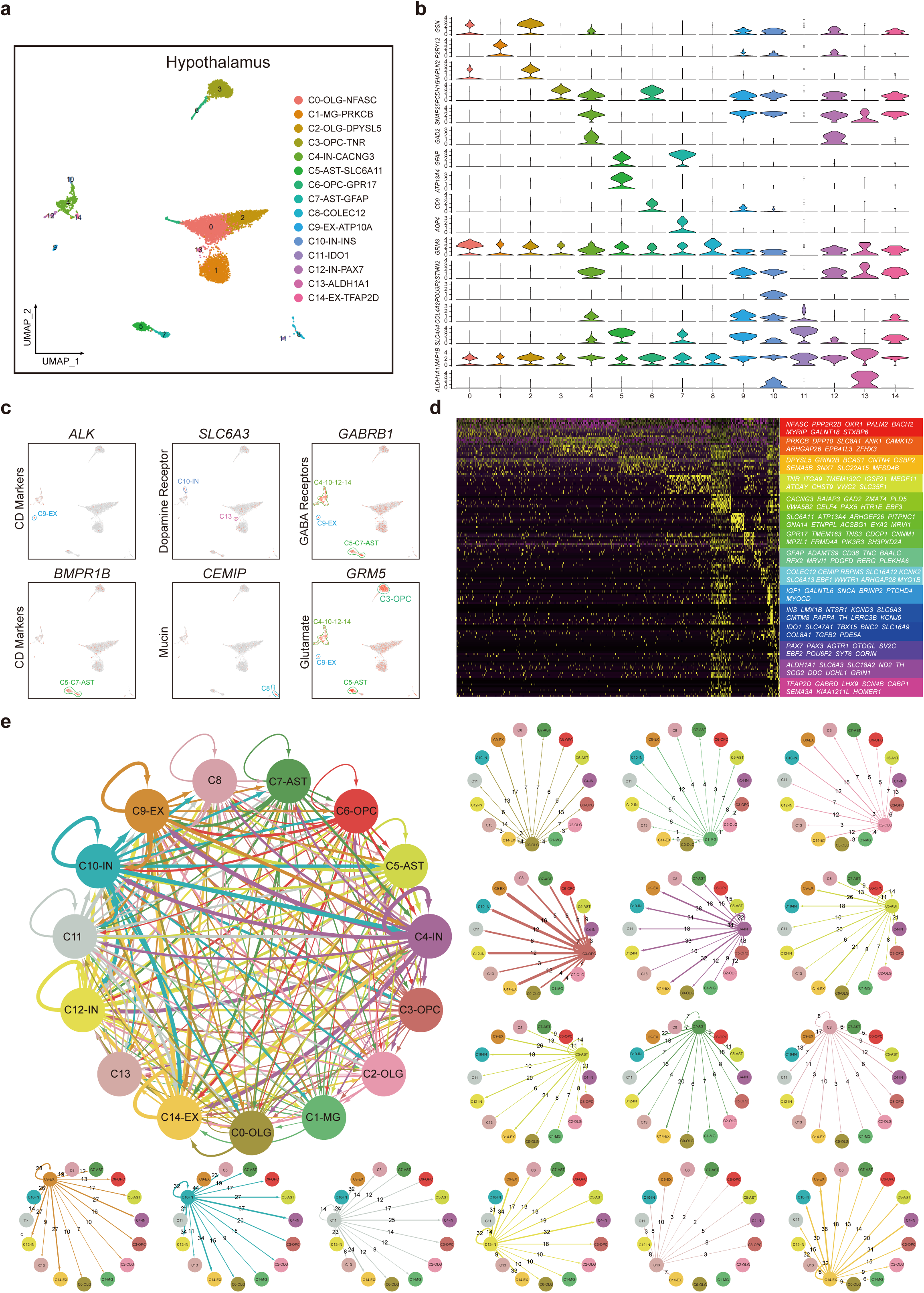

Transcriptomic profiles of receptors and ligands at single cell resolution could help to construct the complex communication network of cell clusters with tissues. Here, we constructed the interaction network of HT cell clusters (see methods), resulting in 2880 pairs of putative interactions mediated by ligands and receptors (Figure 2e). Generally, IN (C4, C10, C12) and EX (C9, C14) interacted most frequently with other clusters. Notably, C1 (MG) interact most frequently with C9 (EX) (Figure S5e, Table S14).

### Cellular heterogeneity and micro-environment in cerebral cortex

Cerebral cortex can be divided into four major regions: FL, PL, OL, TL and each of them was confered a series of crucial biological functionalities. Here we performed unsupervised clustering and cell type assignment separately for each cerebral cortex lobe.

FL principally realizes cognitive skills that play fundamental and critical roles. However, transcriptome at single cell resolution of FL has not been achieved in pig. We identified 12 clusters by clustering 8812 cells of frontal lobe (Figure S2a), and all the 12 clusters were mainly annotated as six major cell types: OLG (C0, C5), MG (C1, C6), OPC (C3), EX (C2, C4, C7, C10), IN (C8) and AST (C9, C11) according to the expression of cell type markers (Figure S2b). In accordance with marker expression, we found cluster specific genes and revealed the molecular marker of FL. Such as *BNC2* and *TCF7L2*, specific expression in C9 (AST), and *TCFL2* is a key transcription factor in the Wnt signaling pathway[66]. Transporter genes *SLC1A2* and *SLC7A11* expression in C11 (AST). *SLC1A2* has been known essential for excitatory neurotransmitter glutamate clear[67] and *SLC7A11* mainly functions in cysteine and glutamate transport systems[68] (FigureS2c). In addition, we found some CD markers that have a specific expression pattern in C4 (EX), C8 (IN), C10 (EX), C11 (AST), such as *CDH2*, *CD47*, and *IGF1R* (Table S7). Intriguingly, the most frequent communications were found within excitatory neurons and identified between excitatory neurons and inhibitory neurons (Figure S2e).

PL incorporates perceptual information from different modalities, including spatial sense, sense of touch, and vision [69]. We generated the PL dataset comprising 6162 cells, resulted in 13 clusters (Figure S3a). Based on reported markers, we identified OLG (C0, C1), OPC (C3, C8), EX (C2, C5, C6, and C9), IN (C10, C11), and AST (C7) cell populations in PL (Figure S3a). We observed that each of the clusters showed distinct gene expression patterns, supporting the validity of the clustering result (Figure S3b). We found that *IL10RB*, a cytokine receptor play an important role in inflammation[70], enriched in C4 (MG). *GLI3* showed a specific expression pattern in C7 (AST), which is an essential TF in sonic hedgehog (Shh) pathway[71](Figure S3c). In addition, there are some markers associated with GABA and glutamate transport system, such as *GABBR2* and *GRIA1* (Table S9). We also performed communication network analysis within parietal lobe. Similar to FL, we found that cellular interplay also enriched in some populations of excitatory neurons (C6 (EX) and C9 (EX)) and inhibitory neurons (C10 (IN)) (Figure S3e).

TL is vital for various functions including the formation of long-term memory, the modulation of auditory, visual sensory input, and the recognition of language[72–74]. Here we provided a dataset containing 3145 individual cells from pig TL. Unsupervised clustering resulted in 12 clusters (Figure S4a), corresponding to major cell types in temporal lobe (Figure S4b). Based on reported markers, we identified OLG (C0), MG (C3), OPC (C1, C4), EX (C2, C6, C8, and C10), IN (C5, C7), and AST (C9) cell populations in parietal lobe. Also, we found the specific expression of several important markers in specific clusters, such as CD marker *IL10RB* in C3 (MG) (Figure S4c). TF encoding gene *TCF7L2* in C1 (OPC), and alpha chain of type XII collagen encoding gene *COL12A1* expressed in C6 (EX) (Figure S4c). Neuron-related markers such as *GABRB1* and *GRM5* showed specific expression patterns in inhibitory neurons and excitatory neurons (Figure S4c). To further dissect the heterogeneity within the temporal lobe, we examined the expression of cluster specific genes and observed clear patterns across clusters (Figure S4d). Cellular ligand-receptor pairs distributed relatively uniformly among clusters compared with other brain regions we examined (Figure S4e).

OL is the center of visual processing and can be divided into several functional visual areas including primary visual cortex and visual association cortex[75]. We sequenced 6829 cells, yielding 16 clusters (Figure S5a). Based on reported markers, we identified OLG (C0), MG (C2), OPC (C5, C13), EX (C1, C3, C4, C6, C8, and C14), and IN (C7, C9, C10, and C12) cell populations in OL (Figure S5b, d). We found that transcription factor encoding gene *RUNX2* was specifically expressed in C2 (MG) (Figure S5c). And *RUNX2* was reported to have an important role in controlling mouse rhythmic behaviors[76]. The transporter gene *ABCA9* was found specifically expressed in the C5 (OPC) (Figure S5c). An appetite regulating gene *NMBR* was found specifically expressed in the C9 (IN) (Figure S5c). CD marker *JAM2* specifically expressed in C11 (Figure S5c), while *CD22* specifically expressed in the C13 (OPC) (Figure S5c). The neural function gene *TACR1* specifically expressed in two inhibitory neuron clusters (C9 (IN) and C11) (Figure S5c). Previous study shows that *TACR1* is associated with several mental health disorders, such as bipolar affective disorder (BPAD), attention-deficit hyperactivity disorder (ADHD), and alcohol dependence syndrome (ADS)[77]. Unlike other cerebral cortex regions, OL showed more extensive and complicated intercellular interactions (Figure S5e). Concretely, neurons, especially C3 (EX) and C4 (EX), broadcasted diverse ligands and showed strong interactions with glia and other neurons.

### Enrichment of disease associated genes in specific cell types

As the primary meat source in Chinese dietary structure, domestic pigs are widely studied to formulate the feed efficiency to control their weights[78,79]. In this section, we aim to gain a deeper understanding of the underlying mechanism of weight regulation in pig hypothalamus, by investigating eating disorders including obesity (OB), Anorexia nervosa (AN), and Bulimia nervosa (BN). We first retrieved diseases associated genes from DisGeNET[80,81] and performed disease enrichment analysis (Methods) to see whether and to what extent, the cell subpopulations contribute to certain diseases. OB, defined as an excess of body fat accumulation for a given height, is the outcome of a gradual energy imbalance between ingestion and expenditure[82]. Among 15 clusters in the hypothalamus, however, no any cluster showed significant enrichment of OB risk genes (Figure 3a). AN, featured as food avoidance and malnutrition, is known as a serious psychiatric disorder[83]. We observed that C2 (OLG), C4 (IN), C9 (OPC), C10 (IN), C12 (IN), and C14 (EX) were significantly associated with AN but to a varying extent (Figure 3b). Notably, we found *DRD2* was distinguishably expressed in C10 (IN) (Figure 3b) and the decreasing expression of *DRD2* could lead to a reduction in food intake. Furthermore, DRD2 could form a heterodimer with ghrelin receptor in neurons in hypothalamus, resulting in the exacerbation of AN[84]. In this regard, we assume that C10 (IN) play a crucial role in AN genesis under DRD2-dependent mechanism. BN is featured with repeated binge eating, following compensatory behaviors[85]. In our hypothalamus dataset, disease enrichment analysis manifested close connection of C4 (EX), C7 (AST), and C10 (IN) with BN (Figure 3c). We noticed that *SLC6A3* displayed specific expression in C10 (IN) (Figure 3c). *SLC6A3* encodes a dopamine transporter, which is responsible for the reuptake of dopamine from the synaptic cleft to presynaptic neurons. Several studies have reported that dopamine plays an important role in the energy intake process, especially when it comes to high-calorie foods[86].

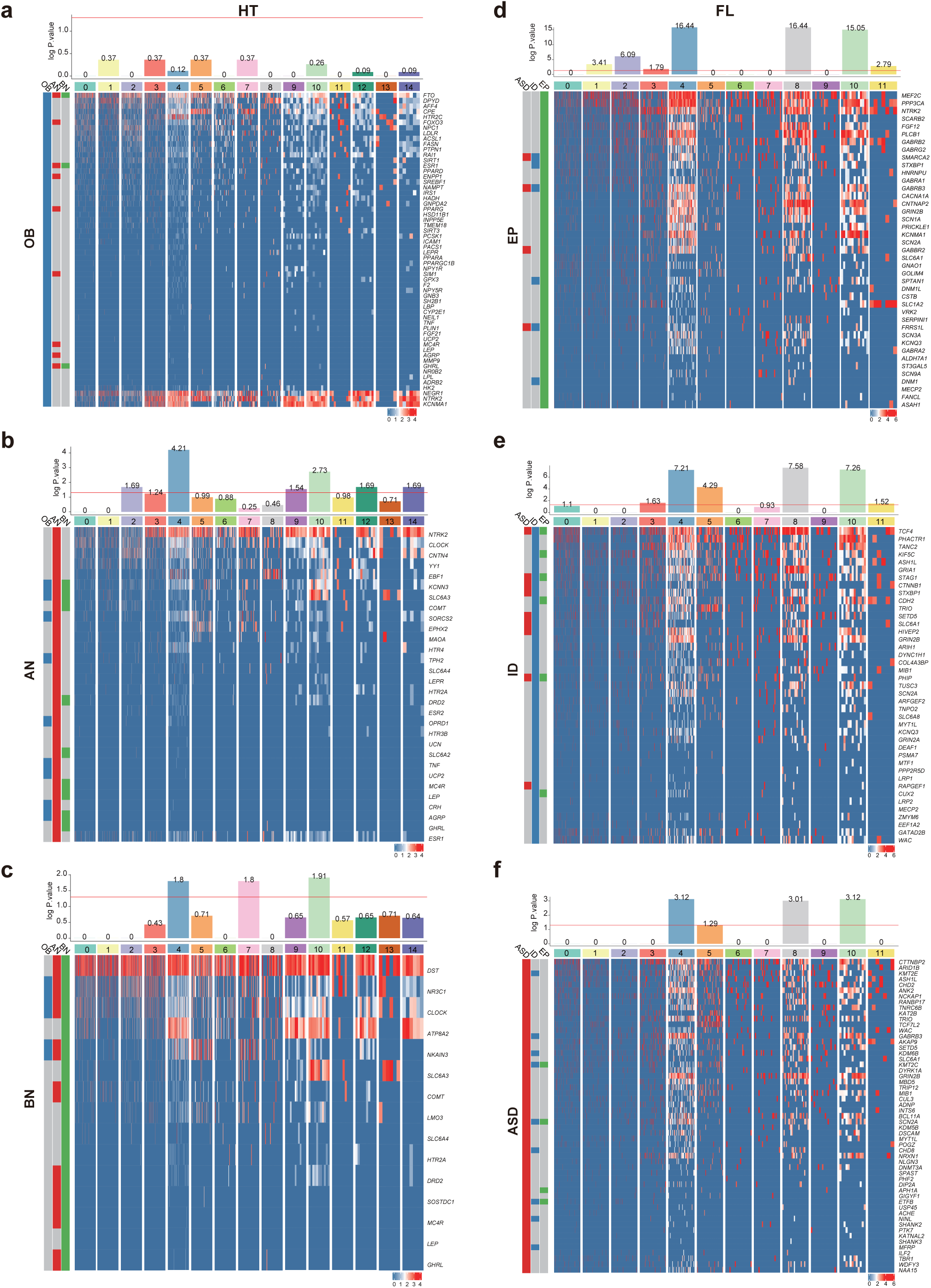

Employing the similar method, we next utilized the comprehensive molecular atlas of pig frontal lobe to identify the cell types where hazardous mutations for neuropsychiatric disease take effect, so as to provide a reference for further researches regarding neuropsychiatric disease mechanisms and circuits. Genes associated with neuropsychiatric diseases, including epilepsy (EP), intellectual disability (ID), and autism spectrum disorder (ASD), has been identified using experimental or gnomic methods in recent years. However, the link between gene mutation and disease progression is still missing. To link neuropsychiatric diseases to specific cell types, we performed enrichment analysis of neuropsychiatric diseases related genes on each cell type of the other four brain regions respectively. In the analysis of FL, we first examined the enrichment of high confidence EP associated genes[87]. Certain neurons subpopulations, including C4 (EX), C8 (IN) and C10 (EX), were found to be highly associated with the occurrence and development of EP and ID, indicating these neuropsychiatric diseases were primarily connected with neuron malfunctioning. Nonetheless, glial cells were also observed to be related to these disease, but of a much lower significance. Concretely, C1 (MG), C3 (OPC), and C11 (AST) were considered to show enriched expression pattern of EP-risk genes (Figure 3d) while C3 (OPC), C5 (OLG), and C11 (AST) specifically expressed a significant part of ID risk genes [88,89] were enriched in (Figure 3e). Although no glial population was significantly linked to ASD, it is worth noting that the enrichment of ASD-risk genes [90] in C5 (OLG) was nearly significant, indicating its possible contribution to ASD (Figure 3f).

### Divergence of brain functions in pig and mouse

Hypothalamus is a small but crucial part of the brain, shared by all vertebrates[91]. According to molecular genetic evidence, it is widely believed that human, rodent, and pig share the same ancestor about 97 million years ago (Figure 4a)[1]. With the question that to what extent the hypothalamus is conserved in the course of evolution, we performed cross-species comparison by integrating hypothalamus dataset of mouse[92] and pig, to figure out the conserved and divergent cell types in the light of evolution. Unsupervised clustering results yielded 23 clusters, with seven neuron subpopulations (C4, C6, C7, C11, C16, C20, C22) and 16 non-neuron clusters (Figure 4b, c). For the purpose of further characterization of the heterogeneity of 23 clusters, we identified conserved cluster specific markers for the two species. All clusters exhibited distinct molecular patterns, providing further evidence for hypothalamus cellular heterogeneity (Figure 4g). *OPALIN*, a central nervous system-specific myelin protein phylogenetically unique to mammals[93] was specifically expressed in C0 (OLG), C3 (OLG), C9 (OLG), and C13 (OLG). *TNR* and *SEMA5A* showed high expression in C1 (OPC) and C12 (OPC), have previously been reported to be detectable in developing oligodendrocytes[94,95]. *P2RY12*, specifically marking microglial cells in rodent and human CNS[96–98], was observed to show distinct expression in C2 (MG) and C14 (MG). We next noticed that clustering result showed a relatively uniform distribution of two datasets (Figure 4b) with cells from the two species in certain clusters aligned well together, indicating little batch effects. In order to better understand the distinguishing features of each species, we next defined species-specific clusters based on contribution index (Methods). Surprisingly we found two cell populations (C2 (OLG) and C20 (EX)) showed enrichment in pig hypothalamus tissue while 10 clusters were defined as shared clusters between mouse and pig. Additionally, we observed that 11 out of 23 clusters were enriched in mouse hypothalamus, most of which were neurons populations (Figure 4e), suggesting that neurons are relatively divergent while glia cells are more conserved between mouse and pig.

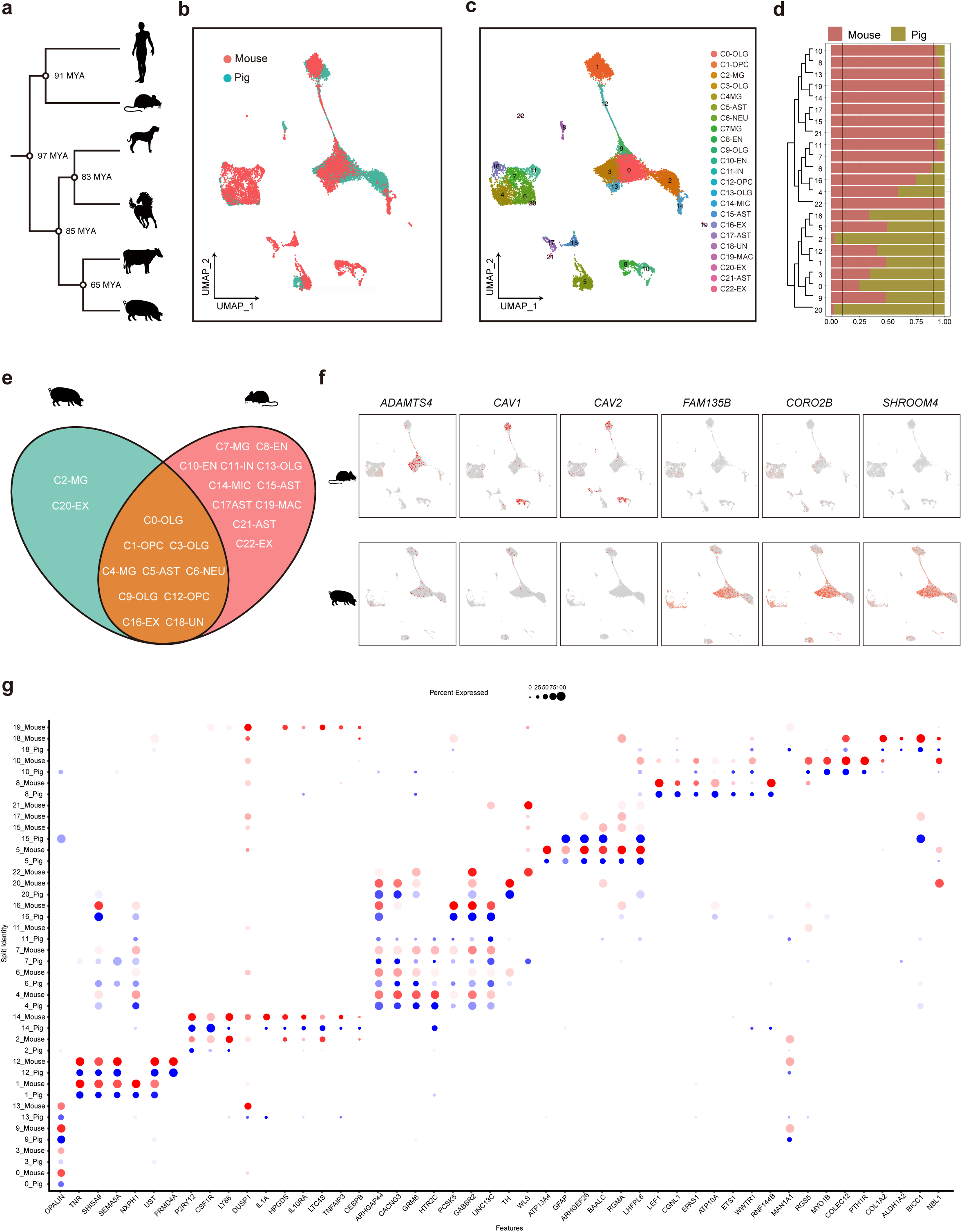

We next sought to discover important genes that were expressed differently between pig and mouse in shared clusters (Figure 4f, Figure S7). We focus on oligodendrocytes due to its pivotal role in evolutionary advancement. Oligodendrocytes, functioning as Schwann cells peripheral nervous system, is responsible for the formation of myelin sheath whose existence is essential for motor, sensory, and higher-order cognitive function[99,100]. Furthermore, oligodendrocytes have proved to be the most abundant glial cells in brain[101]. Firstly in C0 (OLG), we found *ADAMTS4*, a protease that controls the degradation of a brain-specific extracellular matrix protein - brevican[102], specifically expressed in mouse cells (Figure 4f). Previously *ADAMTS4* has been reported to be a marker of mature oligodendrocytes in mouse brain and may promote the generation of myelin sheath[103]. Additionally, *TMEM125* and *TSPAN2*, both known to contribute to neuron myelination in mouse or rat [104,105], also displayed a similar expression pattern as *ADAMTS4* (Figure S7). The scarce expression of these genes (*ADAMTS4*, *TMEM125*, and *TSPAN2*) in pig oligodendrocytes suggested that they may play a more critical role in mouse oligodendrocytes than in pig and the myelination process in pig may be driven by other genes. We also noticed a series of genes, including *PLS1*, *PLIN3*, *ARPC1B*, *ARSG* and so on, that have no reported function in oligodendrocytes evolution but showed distinct expression patterns between mouse and pig (Figure S7). These genes may need further researches to uncover their unknown mechanism of the differences. In C1 (OLG), *CAV1* and *CAV2* were extensively expressed in mouse cells with only sporadic cells expressing *CAV1* and *CAV2* in pig (Figure 4f). CAV1 can heterodimerize with CAV2 to drive the formation of caveolae which can facilitate the high-fidelity neuronal intracellular signaling[106] and modulate the differentiation and regeneration of oligodendrocytes[107]. Besides, *CAV1* is also a primary marker of caveolae in endothelial cells[108], thus explaining the expression of *CAV1* in C8 and C10. On top of that, we observed *C1QL2* exhibit high and specific expression in mouse cells while it was deprived of pig cells (Figure S7). *C1QL2* is a secreted protein[109] that may be engaged in the regulation of the number of excitatory synapses in mouse hippocampus[110]. This may suggest the unknown function of *C1QL2* in hypothalamus and highlight the pivotal role of *C1QL2* in mouse than in pig.

Meanwhile, we also noticed numbers of genes were specifically expressed in pig cells. In C0 (OLG) and C3 (OLG), the expression of *FAM135B* was high and vast in the pig cells, but things are not true in the mouse cells (Figure 4f). In the previous report, scientists found the *FAM135B* may play an important role in the process of Spinal and bulbar muscular atrophy (SBMA)[111]. The survival of normal spinal motor neurons (sMN) derived by iPSC was reduced, after the knockdown of the *FAM135B* gene, and neurite defects were also detected in these cells[111]. It indicated that the *FAM135B* gene may play a more important role in pig sMN growth and survival than in mouse. Besides, *CORO2B* which was known to play critical roles in reorganizing the structure of neuronal actin[112], also exhibits a similar expression spectrum. The higher and more specific expression of *CORO2B* in pig cells than in mouse cells (Figure 4f) may suggest the significance of *CORO2B* in the cell structure organizing of pig oligodendrocytes. *SHROOM4*, which takes part in the process of cytoskeletal organization, was reported critical for human brain structures and survival of some specific neuronal cell types[113]. The deletion of the *SHROOM4* gene was believed may contribute to severe psychomotor retardation and Dent disease[114]. We observed the *SHROOM4* gene was highly and specifically expressed in the pig cells of C0 (OLG) and C3 (OLG), while deprived in the mouse cells (Figure 4f). Implying *SHROOM4* may be more important for the structure of pig oligodendrocytes than mouse by influencing the cytoskeletal organization. Other than those genes, there are also a series of genes showing the same expression pattern as *FAM135B*, *CORO2B*, and *SHROOM4*. Such as *TGFBR3*, *ZDHHC14*, and *MYO1E*, they all show highly and specific expression in pig cells other than mouse cells (Figure S7), indicating their significant roles in the pig brain oligodendrocytes.

## Discussion

Pig has been adopted as a model animal for regenerative medicine research in recent years due to its relatively quick development time and suitable body size. As a putative donor for organ transplantation, the development of pig has been studied widely using genomic and experimental methods. Because of its incredible complexity, the brain remain to be the most mysterious organ to be explored. In this study, we surveyed the pig brain using single cell RNA sequencing followed by comprehensive bioinformatic analysis. The major cell types and related molecular features were identified using unsupervised methods based on single cell transcriptome data. The 32250 cells derived from OL (6829 cells), FL (8812 cells), PL (6162), TL (7302 cells), and HT (3145 cells) were classified into 21 clusters. At spatial level, we identified clusters enriched or depleted in five brain regions, reflecting the cellular composition difference of each brain region. Furthermore, we performed unsupervised clustering for each brain region and identified molecular signatures of TFs, CD marker, ligands, and receptors for OL, FL, PL, TL, and HT, thus providing a comprehensive cellular and molecular taxonomy for pig brain, which represent an valuable resources for understanding pig brain functions at cellular and molecular levels.

Neuropathic diseases such as OB, AN, BN, EP, ID, and ASD have been studied in human brain and a significant proportion of genes have been proposed to be closely related to these diseases. However, there remains a huge gap between gene dysfunction and organ malfunction. As the basic unit of biological functions, cells are crucial for gene expression and function. Several attempts have been made to link human neural diseases to specific cell types. In this study, we systematically assayed six diseases and found the cell types most closely associated with each disease. Overall, we found that different diseases generally linked to a few specific cell types. Our study linked pathology genes to cell types, thus bridging the gap between gene mutation and disease occurrence, which could provide novel insights in understanding disease at cell level. Those cell types related to certain diseases could probably represent targets for medical intervention, which might throw light upon precise medicine based on genomic editing and drug therapy.

The development of organism should be interpreted under evolutionary view. Comparative genomic and transcriptomic studies revealed DNA elements or gene expression modules in different species, which helped us understand species evolution and adaptation at molecular level. The development of scRNAseq brought species-comparison to single cell resolution. The evolution of the neural system in different species reflected their adaptation of its unique environment. The comparison between human and mouse midbrain revealed the species specific features of cell proliferation and developmental timing[31]. The comparative study of monkey and retina reflected the divergence of retina cell types[115]. In this study, we compared the cell diversity and molecular circuits between the hypothalamus of pig and mouse. We found that C0 (OLG), C1 (OPC), C3 (OLG), C4 (NEU), C5 (AST), C6 (NEU), C9 (OLG), C12 (OPC), C16 (EX), and C18 (UN) were shared by mouse and pig hypothalamus. Pig-specific clusters included C2 (MG) and C20 (EX), probably contributing to pig specific biologic functions. Mouse hypothalamus contains more species-specific clusters, most of which were neuron subpopulations, suggesting a more conserved role of glia cells in evolution. Taking a step further, we observed that *ADAMTS4*, *TMEM125*, *TSPAN2* and other genes that are critical in the process of myelination, showed distinctly high expression in mouse oligodendrocytes, indicating the divergent regulation system of myelin sheath generation in pig and mouse.

Taken together, for the first time, we generated a comprehensive cellular atlas for the brain of the domestic pig, an important species of great value in regenerative medicine, agriculture, and evolution. We investigated the composition and molecular features of each cell type and performed spatial and species comparison. Our study revealed divergent cell types at spatial and species levels, as well as conserved cell types with divergent molecular features, which could be invaluable for understanding pig brain development, species specific adaptation and applications to precision medicine.

## Material and methods

### Ethics Statement

The study was approved by the Institutional Review Boards on Ethics Committee of Huazhong Agriculture University. All procedures were conducted following the guidelines of animal experimental committee of Huazhong Agriculture University.

### Sample Collection and Nuclei Extraction

Temporal lobe(TL), frontal lobe(FL), parital lobe(PL), hypothalamus (HT), and occipital lobe(OL) were carefully dissected from a 3 months old male domestic pig following the ethics guidance. Dissected tissues were washed with cold PBS and quickly frozen in liquid nitrogen, then stored in a −80 freezer until use. To start the library construction, tissues were thawed, then cut into small pieces and put into 1.5ml tissue homogenate containing 30mM Cacl_2_, 18mM Mg(Ac)_2_, 60mM Tris-Hcl (pH 7.8), 320mM sucrose, 0.1%NP-40 and 0.1mM EDTA. Tissue homogenate was transferred to 2ml Dounce homogenizer and stroked with the loose pestle for 15 times, followed by 15 times of tight pestle on the ice. The nuclear extraction was filtered with the 40um strainer and spin down at the speed of 500g for 10 min at 4 degrees to carefully discard the supernatant and resuspended with PBS containing 0.1% BSA and 20U/ul RNase Inhibitor prepared for 10x Genomics 3’ library construction.

### Single Cell RNA-seq Library Construction and Sequencing

Libraries were constructed using 10XGenomics V2 kit (PN-120237) following the manufacturer’s instructions and libraries’ conversion were performed using the MGIEasy Universal DNA Library Preparation Reagent Kit and sequenced on BGISEQ-500.

### Pre-processing and Quality Control of Single Cell RNA-seq Data

Cell Ranger 2.0.0 (10X Genomics) was used to process raw sequencing data. Next, Seurat[116] was used for selecting variable genes, dimension reduction, clustering and marker gene identification. Briefly, the top 20 PCs were used for cell clusters and genes with P value of 0.01 and FC of 0.25 were considered as cluster specific genes.

### GO Term and KEGG Pathway Enrichment Analysis

ClusterProfiler R package[117] was used for enrichment analysis and the BH method was employed for multiple test correction. GO terms with a P value lower than 0.01 were considered as significantly enriched.

### Construction of cellular communication network

The construction of cellular communication network of each brain region cells clusters were performed by using our previously described method[118].

### Disease gene enrichment analysis

ASD, ID, EP, OB, AN, and BN high risk gene list were retrieved from DisGeNET[80,81]. We assumed the overlap of cluster-specific genes and disease risk geneset comforn to hypergeometric distribution. enrichment significance was calculated using statistical test, in our case, hypergeometric test. P vlaue is corrected using BH method and 0.05 is used as threshold of significance.

### Cross-species Comparison

Single cell RNA-seq dataset of mouse hypothalamus was downloaded from GSE87544[92]. We first merge expression matrix of the two species (pig and mouse) together based on the intersect of the detected homologous genes. Next, we performed expression matrix preprocessing separately for the two species using Seurat, followed by the integration of two datasets utilizing functions in Seurat. The same filter parameters were chosen as figure 1a. Resolution was set to 1 to yield 23 cell clusters. To determine species-specific clusters, we defined the contribution index *SCI*:

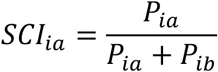

Where *a*, *b* stands for two different species respectively and *P*_*ia*_ denotes the proportion of cells in cluster *i* from species *a* with respect to the total cell number of species *a*. When *SCI*_*ia*_ is greater than 0.1, we defined cluster *i* as species *a* specific cluster.

## Acknowledgement

Dongsheng Chen is supported by China Postdoctoral Science Foundation (grant number 2017M622795). This work was financially supported by Science, Technology and Innovation Commission of Shenzhen Municipality under grant number JCYJ20180507183628543 and the Fundamental Research Funds for the Central Universities [grant numbers:2662018PY025, 2662017PY105]. We are thankful to the production team of China National GeneBank, Shenzhen, China.

## Authors Contributions

G.C., Z.M.L., F.L., D.S.C., F.C., J.X.D conceived and designed the project; D.S.C., J.C.Z. wrote and revised the manuscript. J.CZ., J.X.Z., Y.C., S.Y.W., X.N.D. were responsible for data QC and analysis; X.M.L. and W.Y.W. participated in tissue dissection, sample processing, library construction and experimental verification; F.Y.W., H.Y.W., J.Y.Q. and Y.T.R. participated in data visualization. P.L., G.T., X.Q., J.K.L. contributed to project discussion and technical support.

## Data Availablility

The data that support the findings of this study have been deposited in the CNSA (https://db.cngb.org/cnsa/) of CNGBdb with accession code CNP0000686.

